# *Tex11* Mutant Mouse Models of Human Azoospermia

**DOI:** 10.64898/2026.02.17.706385

**Authors:** Georgia Rae Atkins, Rachel L. Hvasta-Gloria, Chatchanan Ausavarungnirun, Christopher R. Pombar, Jimmaline J. Hardy, Meena Sukhwani, Emily P. Barnard, Nijole Pollock, Michelle Malizio, Yi Sheng, Miguel A. Brieño-Enríquez, Carlos Castro, Tianjiao Chu, Alexander N. Yatsenko, Kyle E. Orwig

**Affiliations:** Department of Obstetrics, Gynecology, and Reproductive Sciences, Magee-Womens Research Institute, University of Pittsburgh School of Medicine, Pittsburgh, PA, United States; Molecular Genetics and Developmental Biology PhD Program, University of Pittsburgh School of Medicine, Pittsburgh, PA, United States; Human Genetics PhD Program, University of Pittsburgh School of Public Health, Pittsburgh, PA, United States; Department of Pathology, University of Pittsburgh School of Medicine, Pittsburgh, PA, United States; Department of Genetics, University of Pittsburgh School of Public Health, Pittsburgh, PA, United States

## Abstract

Non-obstructive azoospermia (NOA) is the absence of sperm in the ejaculate due to spermatogenic failure. Fifty percent of NOA cases are unexplained but may arise from unidentified genetic mutations. Variants in *TEX11* have been identified in men with NOA; and *Tex11* knockout in mice causes NOA. Here we attempt to validate three *TEX11* variants discovered in NOA patients by knocking them into the orthologous region of the mouse genome using CRISPR/Cas9 gene editing. Compared to wild type (144.2 ± 9.87 mg; 1.7 ± 0.5 million sperm/cauda epididymis 4.8 ± 1.3 pups/breeding), *Tex11D* mice (frameshift mutation) had reduced testis weight (28.33 ± 1.16 mg); no sperm in the epididymis; and were infertile with a maturation arrest testicular phenotype. We did not observe any spermatogenesis or fertility defects *Tex11A* mice (missense mutation). *Tex11L* mice had reduced testis weight (87.5 ± 14.79 mg) and epididymal sperm counts (0.33±0.13 million/cauda epididymis) but an incompletely penetrant infertility phenotype (5.4 ± 1.13 pups/breeding) with one third of mice being infertile. Infertile *Tex11L* mice also had a distinct epididymal phenotype with reduced sperm density in the caput and no sperm in the cauda, which was filled with amorphous material.

## Introduction

Infertility is a global public health problem, impacting approximately 17.5% of couples worldwide (World Health 2023). About half of these cases can be attributed to a male factor, which may be caused by lifestyle choices, environmental exposures, occupational hazards, clinically defined syndromes, or genetic mutations (Huang et al. 2023). The most severe form of male infertility is azoospermia, impacting one percent of men in the global population and defined as the absence of sperm in the ejaculate (Cioppi et al. 2021). Azoospermia can be categorized as obstructive or non-obstructive (Cioppi et al. 2021). Obstructive azoospermia (OA) is a physical blockage of sperm passage somewhere within the male reproductive tract (Cioppi et al. 2021). Non-Obstructive Azoospermia (NOA) is spermatogenic failure (Cioppi et al. 2021). NOA is associated the worst clinical outcomes, leaving many with no option of having a biologically related child (Gudeloglu and Parekattil 2013; Agarwal et al. 2015; Esteves 2015; Cioppi et al. 2021). Additionally, 75% of NOA cases are diagnosed as idiopathic, meaning the origin cannot be identified through a standard clinical workup (Yang et al. 2015; Kothandaraman et al. 2016).

Half of infertility is believed to be genetic in origin and several genetic variants have been linked to idiopathic NOA (Zorrilla and Yatsenko 2013; Kasak and Laan 2021). Many of these defects are important for spermatogenesis, meiosis, or DNA repair, including *TEX11/14/15, NR5A1, SYCP1/3, DMRT1, MCM8,* and *MEI1,* (reviewed in (Walsh et al. 2009; Hwang et al. 2010; Massart et al. 2012; Agarwal et al. 2015; Krausz and Riera-Escamilla 2018; Ghieh et al. 2019)). Yatsenko and colleagues reported that *TEX11* mutations are common, occurring in 2.4% of a NOA patient population and 15% in those NOA patients with confirmed maturation arrest by histological analysis (Yatsenko et al. 2015). Several other groups have identified additional *TEX11* variants in NOA patients (Table 1).

**Table 1.** *TEX11* variants identified in NOA patients. Table Note: All variants converted to Isoform 2 transcript position (NM_031276). MAF from gnomAD v.4.0.0. (Chen et al., 2024). Pub. Ref. ID = patient identifier in respective publications. Only pathogenic or variants of unknown significance (VUS) are listed. Likely benign or benign reported variants were omitted.

| Pub. Ref. ID | Diagnosis | Nucleotide Change | Exon | AA Change | MAF | ACMG |
| --- | --- | --- | --- | --- | --- | --- |
| **Song et al., 2023** |  |  |  |  |  |  |
| Family 1 | NOA | c.2575G>A | ex29 | p.G859R | N/A | VUS |
| Family 2 | NOA | c.313C>T | ex5 | p.R105* | N/A | Pathogenic |
| **Ghieh et al., 2022** |  |  |  |  |  |  |
| P20 | NOA | c.1483G>A | ex18 | p.A495T | N/A | VUS |
| **Tang et al., 2022** |  |  |  |  |  |  |
| NOA39 | NOA | c.466A>G | ex7 | p.M156V | 0.0003065 | VUS |
| NOA49 | NOA | c.559_560del | ex8 | p.M187fs | N/A | Pathogenic |
| **Ji et al., 2021** |  |  |  |  |  |  |
| P5648 | NOA | c.1751 + 2T > G | Int20 | p.584K spl d | N/A | Pathogenic |
| P6825 | NOA | c.1381-1C > T | Int16 | p.461A spl d | N/A | Pathogenic |
| P7583 | NOA | c.2653G>T | ex29 | p.W856C | N/A | Pathogenic |
| P5048 | NOA | c.1006G > T | ex11 | p.E336* | N/A | Pathogenic |
| P8122 | NOA | c.1209dupA | ex15 | p.N403fs*10 | N/A | Pathogenic |
| P8251 | NOA | c.253delG | ex4 | p.V85fs*5 | N/A | Pathogenic |
| P9225 | NOA | c.812delA | ex11 | p.K271fs*5 | N/A | Pathogenic |
| **Yu et al., 2021** |  |  |  |  |  |  |
| N/A | NOA | c.151_154del | ex3 | p.D51fs | N/A | Pathogenic |
| **An et al., 2021** |  |  |  |  |  |  |
| A2799 | NOA | c.2240C>A | ex26 | p.S747* | N/A | VUS |
| A4999 | NOA | c.1337G>T | ex16 | p.R446M | N/A | VUS |
| A961 | NOA | c.466A>G | ex7 | p.M156V | 0.0003065 | VUS |
| A2153 | NOA | c.1246C>T | ex16 | p.Q416* | N/A | Pathogenic |
| **Krausz et al., 2020** |  |  |  |  |  |  |
| 09-297 | NOA | c.39_606del | ex3-8 | p.13del189aa | N/A | Pathogenic |
| **Sha et al., 2018** |  |  |  |  |  |  |
| Patient 1, 2 | NOA | c.2653G>T | ex29 | p.W856C | N/A | Pathogenic |
| **Nakamura et al., 2017** |  |  |  |  |  |  |
| Patient 3 | NOA | c.466A>G | ex7 | p.M156V | 0.0003065 | VUS |
| **Yang et al., 2015** |  |  |  |  |  |  |
| WHT3150 | NOA | c.349T→A | ex6 | p.W117R | 0.000001836 | VUS |
| WHT3500 | NOA | c.515A→G | ex7 | p.Q172R | N/A | VUS |
| WHT3759 | NOA | c.1258Ins (TT) | ex16 | p.D435fs | N/A | Pathogenic |
| **Yatsenko et al., 2015** |  |  |  |  |  |  |
| 1, 3 | NOA | c.607del237bp | ex9-11 | p.203del79aa | N/A | Pathogenic |
| 2 | NOA | c.466A>G | ex7 | p.M156V | 0.0003065 | VUS |
| 4 | NOA | c.1793+1G>C | Int21 | p.R597spl d | N/A | Pathogenic |
| 5 | NOA | c.747+1G→A | Int10 | p.L249spl d | N/A | Pathogenic |
| 7 | NOA | c.2047G→A | ex24 | p.A683T | 0.0002910 | VUS |

Yang and colleagues reported that *Tex11* interacts with *Sycp2*, a component of the synaptonemal complex lateral elements crucial for homologous recombination during meiosis (Yang et al. 2008). *Tex11*-deficient spermatocytes exhibit loss of synapsis and crossover, leading to meiotic arrest at the pachytene stage and infertility (Adelman and Petrini 2008; Yang et al. 2008). We modeled three different human NOA-associated TEX11 variants in mice to validate the genotype/phenotype association. Tex11D is a frameshift mutation (p.D435fs); Tex11A is a missense mutation (p.A683T); and Tex11L is a splice site mutation (p.L249). Reproductive phenotyping in mouse models will test the overarching hypothesis that these NOA patient-associated variants in *TEX11* are the cause of infertility.

## Results

### Fertility/infertility phenotypes of Tex11 variants

TEX11 is well conserved (75% nucleotide identity; 56% amino acid identity) between humans and mice with identical conserved domains (Fig. 1A). Each of the human mutations were modeled in the orthologous regions of the mouse *Tex11* gene. The frameshift mutation at p.D435fs (*Tex11D*) is within the TPR-like domain and is designated “Pathogenic” by the American College of Medical Genetics (ACMG) guidelines for variant interpretation (Table 1). The missense mutation at p.A683T (*Tex11A*) is a “Variant of Unknown Significance (VUS)” and not within a currently known functional domain. However, the substitution of Threonine for Alanine may introduce steric hindrance into the predicted tightly packed helix-turn-helix region of the protein. The splice site mutation at p.L249spl (*Tex11L*) is predicted “Pathogenic” by ACMG and is in the meiosis specific Spo22/ZIP4/Tex11 domain. Since each of these variants are in different domains of *Tex11* and cause different kinds of mutations with different predicted pathogenicity, we hypothesized that they would have different impacts on fertility.

**Figure 1.**
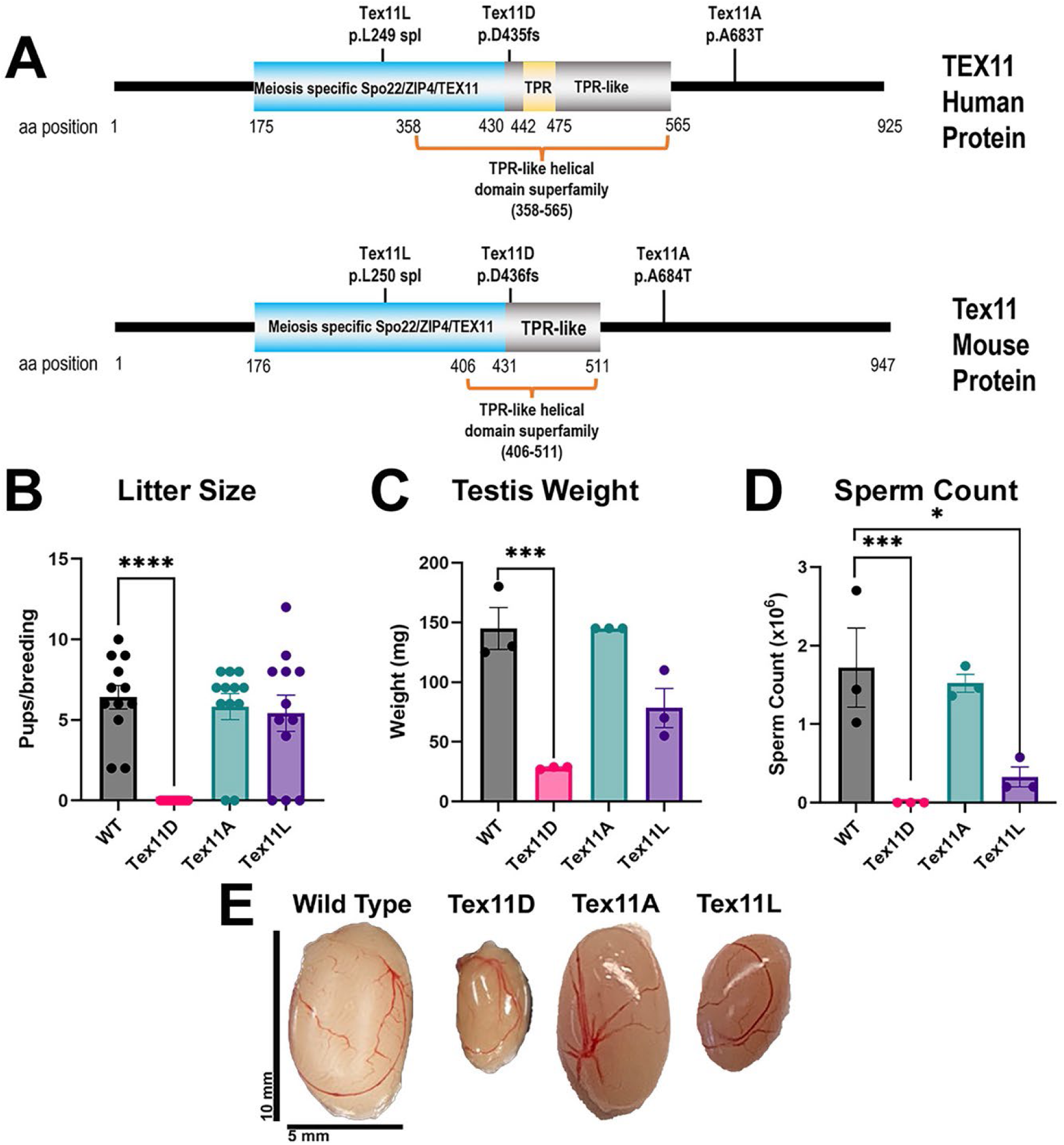
*Tex11* mice modeling different human *TEX11* variants demonstrate a variable impact on fertility. (*A*) Locations of *TEX11* variants identified in patients with NOA on human TEX11 and corresponding locations on the mouse protein. (*B*) The number of pups per breeding with wild type females. (n=3 for all lines except n=5 for *Tex11D)* ****p< 0.0001 (*C*) Testis weights of each line in mg (n=3/line age 4-6 months) ***p=0.0037. (*D*) Sperm count calculated from sperm retrieved in cauda epididymis and reported as million/cauda (n=3/line) *p=0.042***p=0.009. (*E*) Brightfield images of testes from each mouse line.

Breeding analyses indicated that *Tex11D* mice were infertile with zero litters produced from ten matings compared to wild type males that produced 12 litters from 12 matings (6.42±0.73 pups/breeding, p<0.0001). *Tex11A* mice produced 10 litters from 12 matings (7.00±0.26 pups/breeding, p=0.46) and *Tex11L* mice produced 9 litters from 12 matings (7.22±0.83 pups/breeding, p=0.43), which were not significantly different than wild type. The testes of *Tex11D* mice (28.33±1.16 mg) were smaller than wild type (145±17.56mg; p=0.011, Fig. 1C and D)), while the testis size of *Tex11A* mice (145±0 mg, p=0.92) and *Tex11L* mice (78.33±16.41 mg, p=0.13) were not significantly different than wild type (Fig. 1C and E).

Furthermore, *Tex11D* had no sperm in the cauda epididymis, while wild type mice had 1.72±0.5 million/cauda epididymis (p=0.009). Epididymal sperm counts in *Tex11A* mice (1.52±0.0.11 million/cauda) were not statistically different than wild type (p=0.89). *Tex11L* mice (0.33±0.13 million/cauda epididymis) had statistically lower sperm counts than wild type (Fig. 1D, p=0.042). Interestingly, one of the *Tex11L* mice produced only one litter, while the other two TEX11L mice produced several litters, suggesting an incompletely penetrant infertility phenotype, which is tested more rigorously below.

### The Tex11D mutation causes NOA with disruptions in meiotic prophase I in mice

To understand the stage of spermatogenesis impacted by the *Tex11* variants, we utilized histological analysis of the testes and caput epididymis (Fig. 2). *Tex11D* mice had VASA+ germ cells but no TP1+ spermatids, suggesting a maturation arrest phenotype (Fig. 2F). As expected, TEX11 protein expression was absent in the *Tex11D* mice, demonstrating that the frameshift mutation leads to complete loss of TEX11 protein (Fig. 2G). The caput epididymis of *Tex11D* mice contained round cells but no sperm (Fig. 2H). Immunohistochemistry revealed that *Tex11D* mouse seminiferous tubules contained SALL4+ undifferentiated spermatogonia, STRA8+ differentiating spermatogonia, and PIWIL1+ spermatocytes, but not TP1+ postmeiotic spermatids (Supplemental Fig. 2). Thus, the *Tex11D* mouse model recapitulated the maturation arrest phenotype observed in the human patient and validated the p.D435fs NOA-associated variant (Yang et al. 2015; Wang et al. 2021).

**Figure 2.**
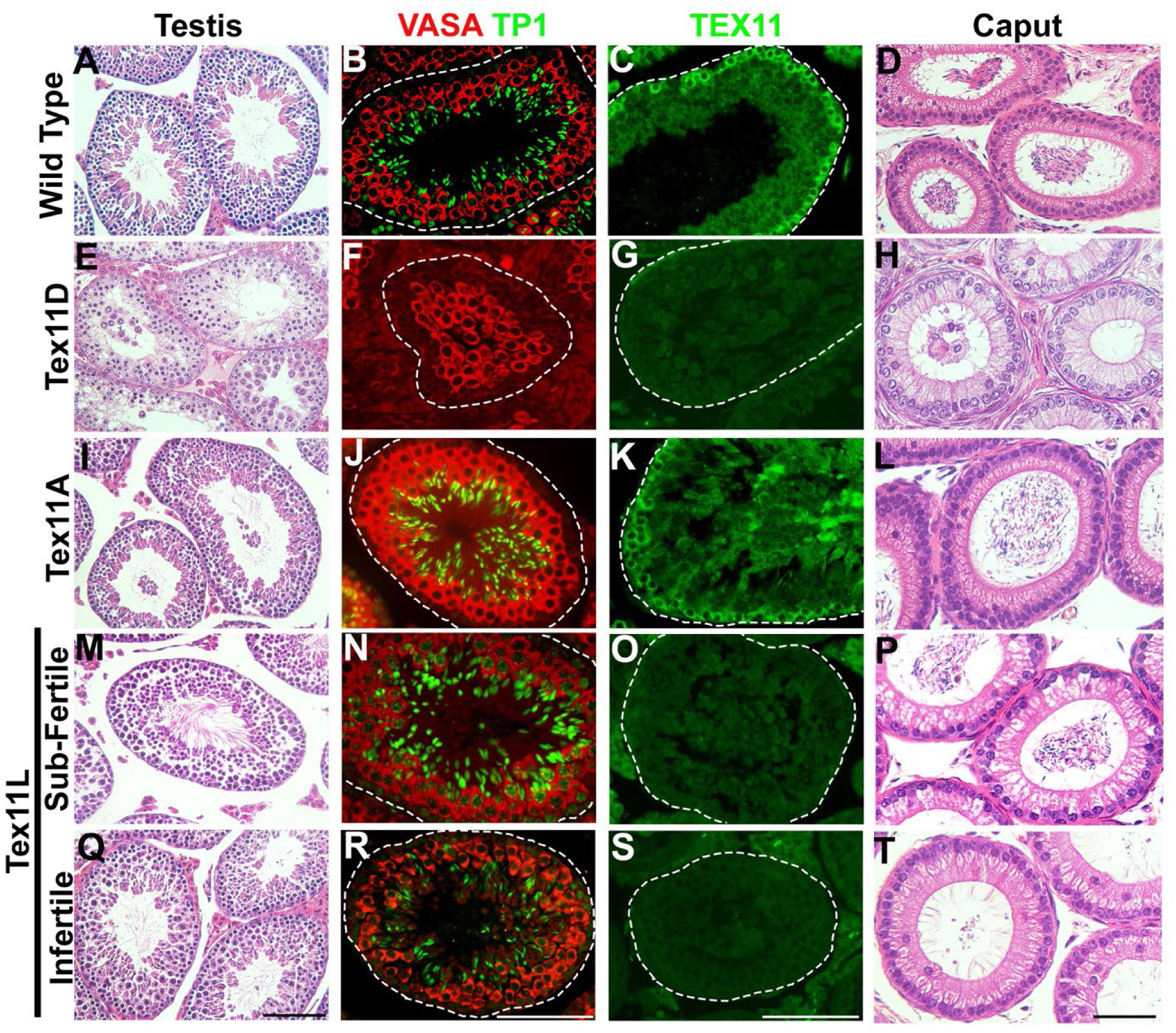
Testicular and epididymal phenotypes of *Tex11* mutant mice. Testis and epididymal histology in (*A-D*) wild type, (*E-H*) *Tex11D*, (*I-L*) *Tex11A*, (*M-P*) *Tex11L* sub-fertile and (*Q-T*) *Tex11L* infertile mice. (*A, E, I, M, Q*) Hematoxylin and eosin staining of testicular cross sections. (*B, F, J, N, R*) Immunofluorescence co-staining of seminiferous tubules with TP1 (green, spermatid marker) and VASA (red, germ cell marker). (*C, G, K, O, S*) TEX11 expression in the testis. (*D, H, L, P, T*) Hematoxylin and eosin staining of caput epididymis cross sections. Scale bar in all images is 100 µm.

Previous analysis revealed that meiotic pairing, synapsis, and recombination defects occur in *Tex11* knockout mice (Yang et al. 2008). To confirm whether our Tex11D mutation replicates the previously observed phenotype, we performed meiotic prophase I spreads. Meiotic prophase progression is characterized by two features: synapsis of homologous chromosomes and the initiation of meiotic recombination through DNA double-strand break (DSB) formation.

We defined the meiotic prophase I substages based on the cytological appearance of synaptonemal complex proteins (SYCP3). We assessed meiotic synapsis progression using antibodies against SYCP3 and SYCP1. We observed that *Tex11D* mice exhibited an increase in synapsis defects showing approximately 48% abnormal compared to only 18% abnormal in control mice (Fig. 3A, B). Meiotic recombination is a precisely regulated process, with DSBs generated throughout the genome in a highly controlled and specific manner. We evaluated DSB occurrence using antibodies against γH2AX and SYCP3, finding that approximately 54% of *Tex11D* spermatocytes show γH2AX retention or mislocalization (flares and bubbles) during pachynema compared to only 20% in controls (Fig. 3C, D). To investigate the cause of the repair defect, we analyzed the spatial and temporal distribution of repair proteins. RAD51 marks sites of double-stranded breaks, serving as an early recombination nodule marker. It first appears during leptonema and mostly resolves as breaks are repaired. RPA binds to single-stranded DNA and appears during leptonema, resolving as meiosis progresses, indicating a transition to a meiotic nodule. Both markers decrease as DSBs are repaired or crossovers occur. In *Tex11D* mutants, there was no difference in the number of RAD51 or RPA foci at leptonema compared to wild type (RAD51 control=67±3.78 vs RAD51 *Tex11D*=60±4.02; RPA control=87±4.98 vs RPA *Tex11D*=98±5.67). However, a significant increase was observed in both foci during zygonema (RAD51 control=127±5.33 vs RAD51 *Tex11D*=146±6.04, p=0.0007; RPA control=202±8.08 vs RPA *Tex11D*=251±8.39, p<0.0001) and pachynema stages (RAD51 control=5±0.48 vs RAD51 *Tex11D*=28±1.18, p<0.001; RPA control=5±0.64 vs RPA *Tex11D*=28±1.37, p=0.003) (Suppl. Fig. 4A,B), indicating a failure to resolve DSBs and progress to a meiotic nodule. Proper repair is essential for meiotic recombination, leading to crossover formation. We evaluated this using the MLH1 crossover marker. The control group averaged 23 MLH1 foci, while *Tex11D* averaged 18 foci (Suppl. Fig. 4C, p<0.0001). Overall, these results suggest that the *Tex11D* mutation disrupts meiotic prophase I, consistent with the findings reported by Yang et al. in *Tex11* knockout mice (Yang et al. 2008).

**Figure 3.**
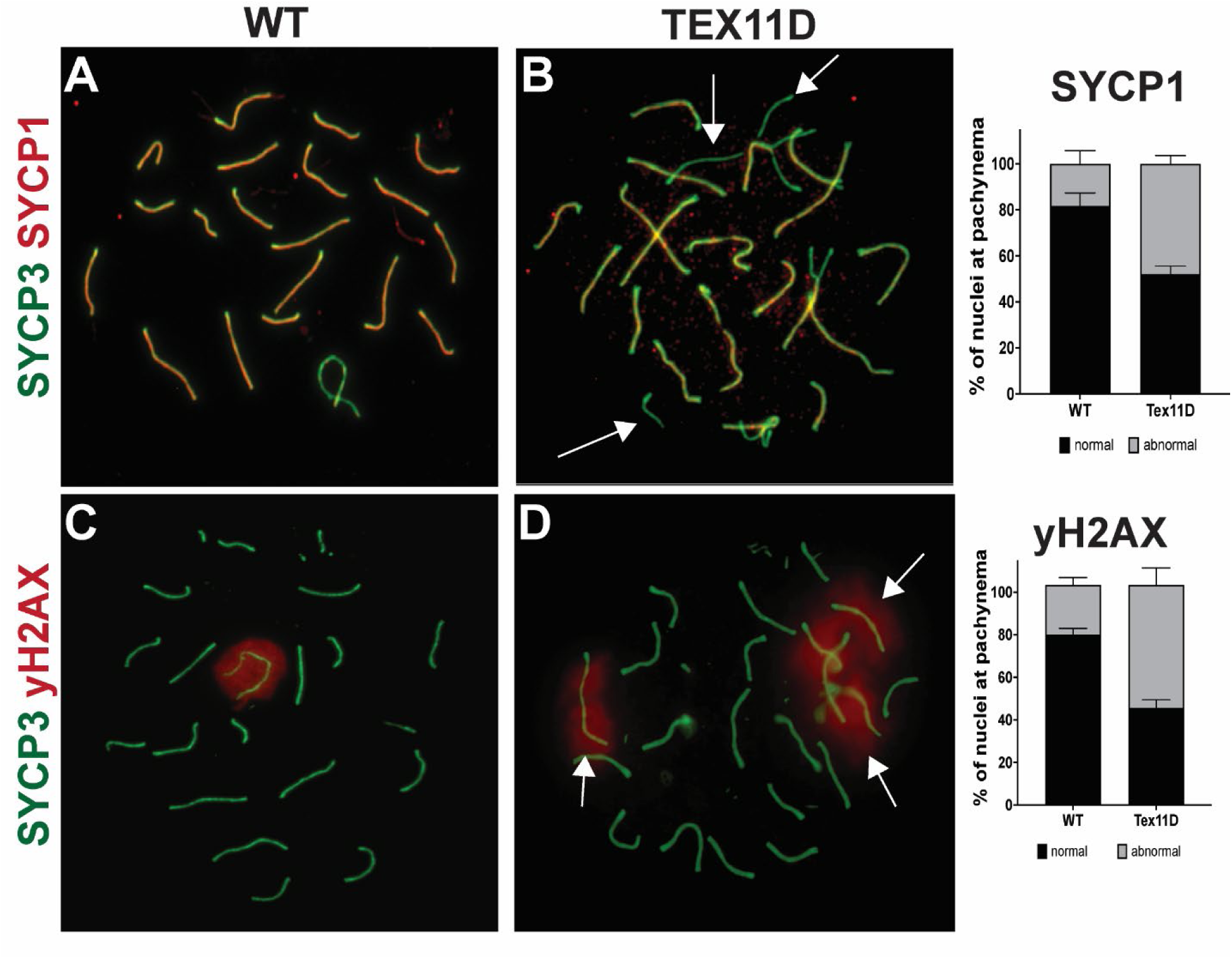
*Tex11D* mice show increased chromosome asynapsis. (*A, B*) Meiotic spreads staining for SYCP3 (green) and SYCP1 (red) to demonstrate chromosome synapsis. Arrows indicate asynapsed chromosomes, which are quantified in graph in the right panel. Bars represent mean ± SEM. (*C, D*) Meiotic spreads staining for SYCP3 (chromosomes, green) and γH2AX bodies (red) in wild type and mutant mice. Arrows show unexpected γH2AX bodies in addition to the region over the sex chromosomes that is typically seen. Quantified in graph on the right. Bars represent mean ± SEM.

### Tex11A mutant mice have normal spermatogenesis

Spermatogenesis in the seminiferous tubules of *Tex11A* mice exhibited normal tubule histology; VASA (germ cells) and TP1 (spermatids) expression were indistinguishable from wild type mice; and sperm were present in the lumen of the caput epididymis (Compare Fig. 2A-D with Fig. 2I-L).

### Tex11L causes variable impact on spermatogenesis

*Tex11L* mice presented with two distinct phenotypes: sub-fertile and infertile. At 12 weeks, two of the *Tex11L* mice produced litters (4 litters in 4 breeding attempts), while the other *Tex11L* mouse only produced one litter in four breeding attempts (Fig. 1B). All *Tex11L* mice had lower sperm counts and testis size compared to wild type mice (Fig. 1C-E). Sub-fertile mice were characterized by decreased testis weights and sperm counts but produced multiple litters, while the infertile mouse had decreased testis weight and sperm count and produced only one litter in four breeding attempts. We predicted that *Tex11L* may cause an incompletely penetrant infertility phenotype, resulting in some tubules with maturation arrest and others with complete spermatogenesis. At 12-weeks, testes from *Tex11L* sub-fertile (n=2) and infertile (n=1) mice exhibited normal seminiferous tubule morphology with VASA+ germ cells and TP1+ spermatids (Fig. 2M-T). Thus, there was no obvious impact of the *Tex11L* splicing mutation on spermatogenesis at 12 weeks. However, no TEX11 protein expression was detected in the sub-fertile or infertile *Tex11L* mutants (Fig. 2O, S). This could be an antibody problem because the antibody used was against the ZIP4H/TEX11 domain and the *Tex11L* splice site mutation is in the middle of that domain. All commercially available antibodies for mouse TEX11 use the ZIP4H/TEX11 domain as the immunogen. It is assumed that there is TEX11 expression in the *Tex11L* mice because sperm were found and TEX11 is required for spermatogenesis. In contrast to testis morphology and immunofluorescent results, the caput epididymis contained sperm in the sub-fertile group but not the infertile group (Fig. 2P, T).

### Tex11L infertile mice have an abnormal cauda epididymis that worsens with age

To confirm the reproducibility of the infertile *Tex11L* phenotype (n=1) and whether the phenotype was age dependent, three *Tex11L* hemizygous males and three wild type littermates were bred continuously for 52 weeks to wild type females (two females/male). Every two to three weeks, breeding females were removed and replaced with two new breeding females. All wild type males (Male 1-3) were fertile, producing 23/25, 18/20 and 11/20 litters per breeding attempt (Fig. 4A). Meanwhile, two hemizygous males were fertile, producing 18/24 and 15/20 successful breeding attempts (Male 4 and 6), while one male had only two litters out of 19 breeding attempts (Male 5) (Fig. 4A). Hemizygous Male 5 had a drastically lower sperm count and testis weight compared to the other two hemizygous males and wild type males, similar to the results for one of the three hemizygous males in Figure 1B-E.

**Figure 4.**
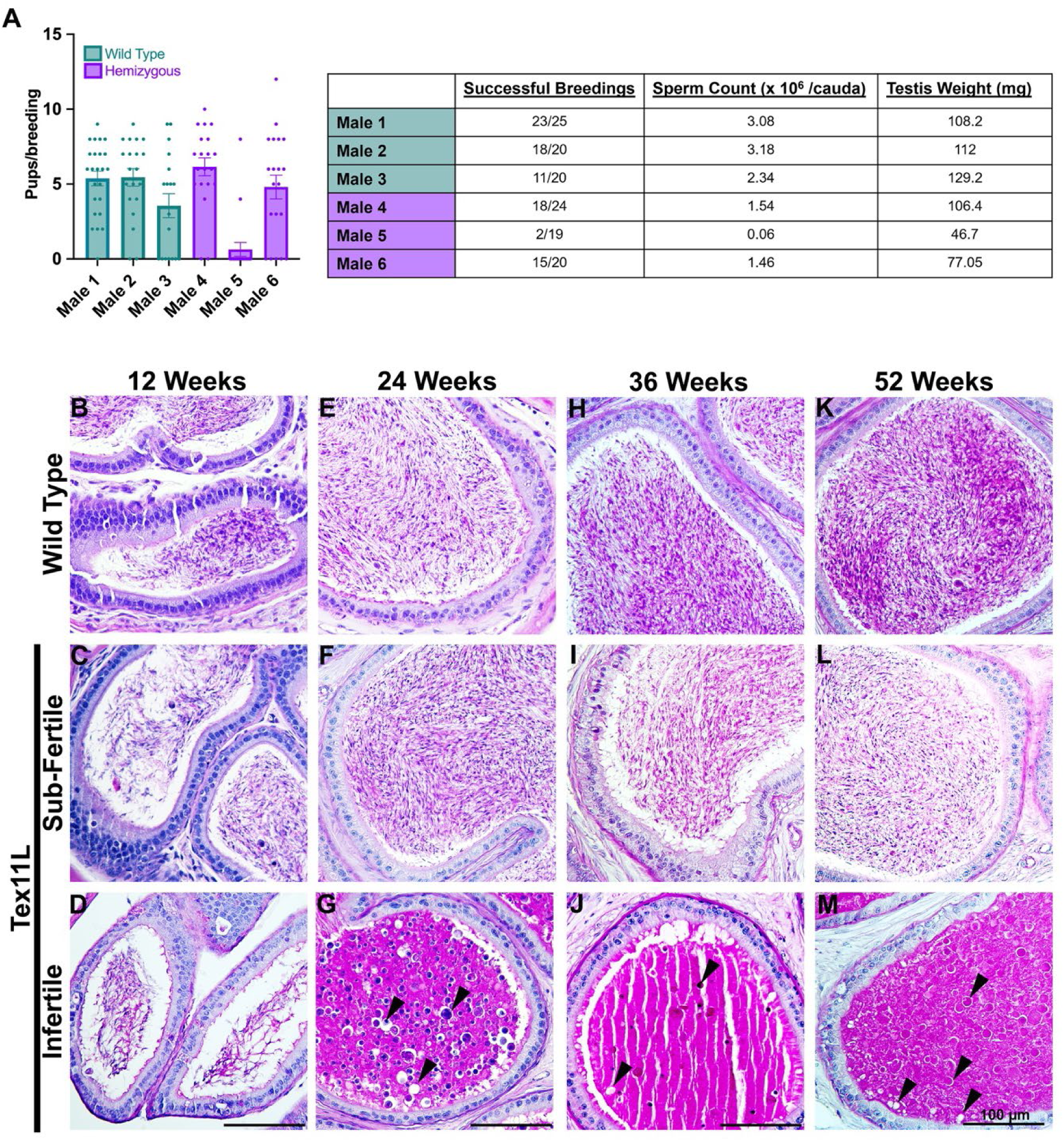
Some Tex11L mice have an infertile phenotype with aberrant morphology in the cauda epididymis beginning at 24 weeks. (*A*) *Tex11L* wild type littermates and hemizygous males bred for 52 weeks show the same phenotype seen at 12 weeks with two males being sub-fertile (produced offspring, but reduced sperm count and testis weight) and one male being infertile. Periodic acid-Schiff stain of wild type, *Tex11L* sub-fertile, and *Tex11L* infertile mice cauda epididymides from 12 (*B-D*), 24 (*E-G*), 36 (*H-J*), and 52 weeks (*K-M*). Wild type (*B, E, H, K*) and *Tex11L* sub-fertile (*C, F, I, L*) mice had similar cauda epididymis phenotypes with abundant sperm in the lumen. Starting at 24 weeks and persisting through 52 weeks, the cauda epididymides in the *Tex11L* infertile mice had an accumulation of macromolecular substance and round bodies in the lumen (*G, J, M*). (Wild type n=3/time point, *Tex11L* sub-fertile n=2/time point, *Tex11L* infertile n=1/time point). Scale bar in all images is 100 µm.

Additionally, *Tex11L* sub-fertile and infertile animals displayed abnormal seminiferous tubule morphology (Suppl. Fig. 4). The sub-fertile animals had a hypospermatogenesis phenotype with some normal seminiferous tubules and some atrophic or disorganized seminiferous tubules. All seminiferous tubules in the infertile *Tex11L* mice exhibited a maturation arrest phenotype at 52 weeks of age.

To further characterize the abnormal testicular morphology and infertility phenotype seen in *Tex11L* infertile mice, mutant testes and epididymides were analyzed at 12, 24, 36 and 52 weeks of age (n=1/timepoint). While sperm were observed in the cauda epididymis at 12 weeks in wild type, *Tex11L* sub-fertile and infertile animals, *Tex11L* infertile mice developed a distinct cauda epididymis phenotype at 24 weeks and beyond that was characterized by an amorphous luminal substance containing vacuoles and sloughed cells (Fig. 4B-M). The phenotype was not apparent at any timepoint in wild type or *Tex11L* sub-fertile mice. We also analyzed the size of seminiferous tubules in wild type and *Tex11L* infertile mice by measuring length x width cross-product of each tubule at each time point. Fifty tubules per mouse were measured and the averages were used for statistical analysis. The average tubule size was significantly decreased at 12, 24, 36, and 52 weeks in *Tex11L* infertile mice compared with wild type (p<0.0001; Fig. 5A). In addition, the localization of TP1+ spermatids in the seminiferous tubules of *Tex11L* infertile mice was abnormal compared with wild type seminiferous tubules, a phenotype that worsened with age. TP1+ signal typically appears as a tight ring of spermatids surrounding the seminiferous tubule lumen (Fig. 5B, D, F, H). The *Tex11L* infertile mice had TP1+ cells sloughed in the lumen as well as a reduced number of TP1+ cells (Fig. 5C, E, G, I).

**Figure 5.**
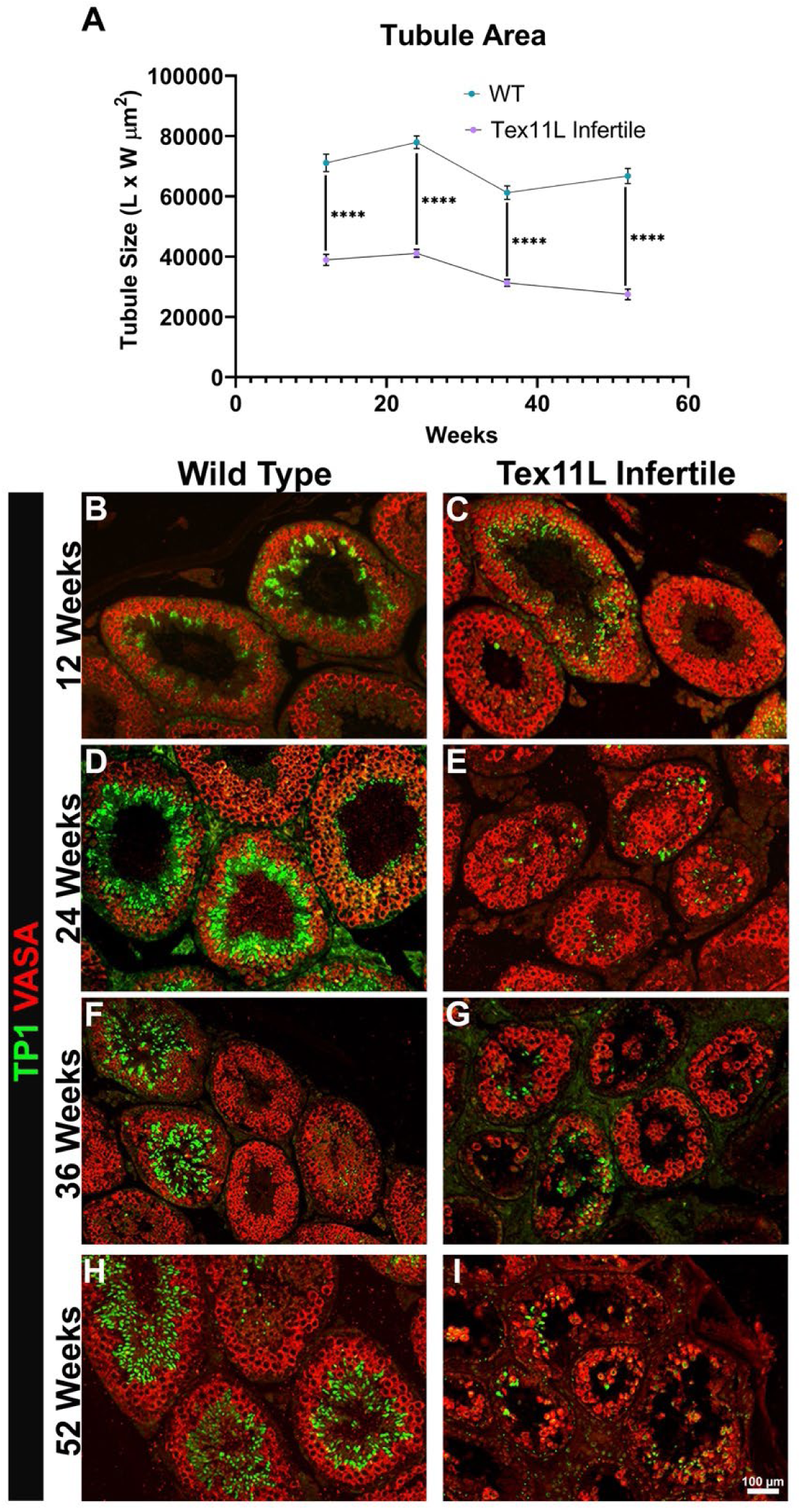
*Tex11L* infertile mice have decreased tubule size and an irregular seminiferous epithelium. (*A*) Tubule size was measured by the length and width of each tubule. Average size is shown on the graph. One mouse for each genotype was analyzed at 12, 24, 36, and 52 weeks and 50 tubules for each mouse were counted and used to calculate the average. ****p<0.0001. Immunohistochemistry of testis tissue with TP1+ spermatids and VASA+ germ cells in wild type and *Tex11L* infertile hemizygous mice at 12 (*B-C*), 24 (*D-E*), 36 (*F-G*), and 52 (*H-I*) weeks. Scale bars=100 µm.

## Discussion

In this study, we created and evaluated three mouse models with human NOA-associated *TEX11* mutations: one frame shift mutation, one missense mutation, and one splicing mutation (Table 2). Histological analysis of the frameshift mutation (*Tex11D*) in mice revealed that germ cell maturation was arrested at the spermatocyte stage, which replicated the phenotype that was previously reported in mice (Wang et al. 2021) and the human patient (Yang et al. 2015). No sperm were observed in the testis or the caput or cauda of the epididymis of *Tex11D* mice and they failed to sire litters after breeding with wild-type females.

**Table 2.**
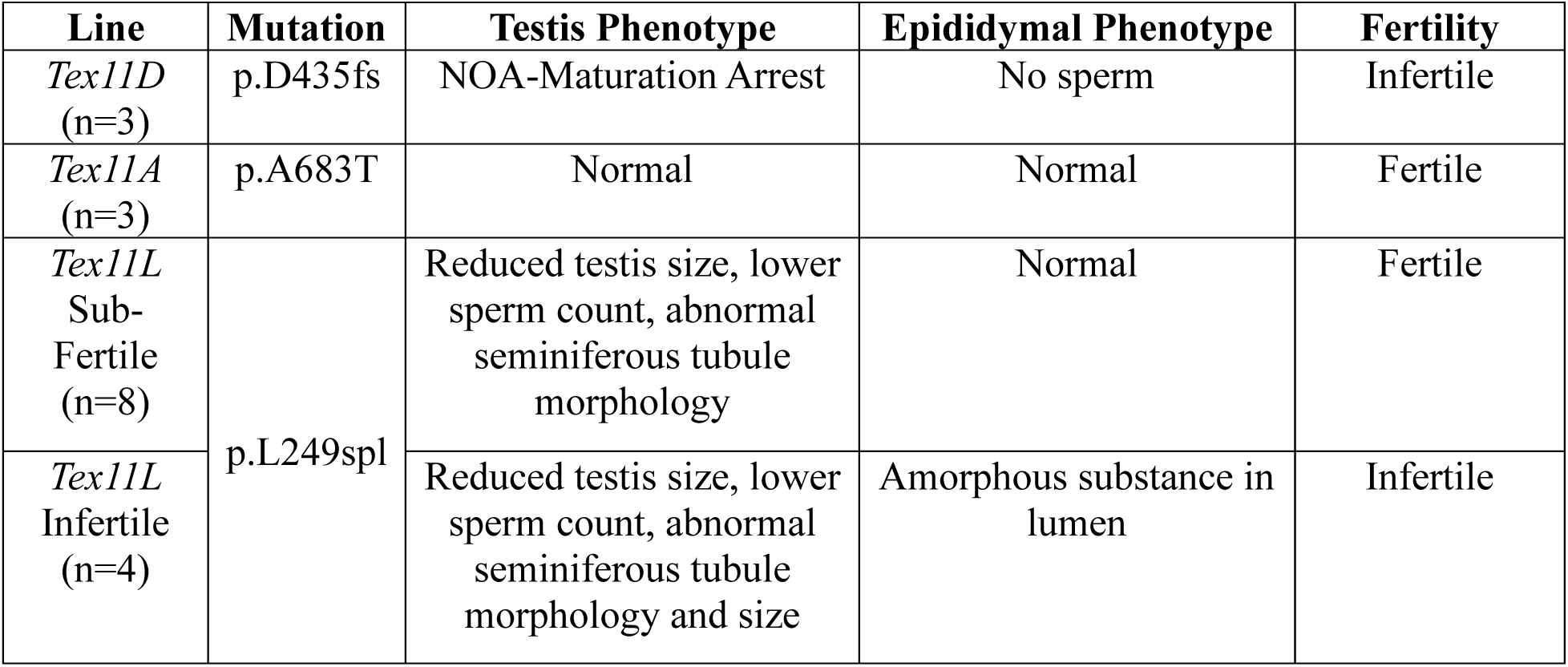
Summary of phenotypes seen in mouse models of NOA-associated *TEX11* variants.

The *Tex11A* mutation occurred in an uncharacterized region of the protein, situated between the TPR-like domain and the C-terminus, featuring densely packed helix-turn-helix motifs. The mutation occurred at the start of the residue 682 loop, which we hypothesized would introduce steric hindrance, thus obstructing the tertiary fold of the protein. However, the *Tex11A* line was fertile with no obvious morphological defects to the testis or epididymis. Previous mouse models of human missense variants in the mouse *Tex11* gene include p.W117R, p.Q172R, and p.V748A knock-ins (Yang et al. 2015). Mice with all three missense mutations were fertile but p.V748A mice had significantly reduced sperm counts and asynapsis in germ cells (Yang et al. 2015).

The *Tex11L* splicing variant occurred within the highly conserved meiosis specific Spo22/ZIP4/Tex11 functional domain, which has been previously shown to be necessary for forming foci between homologous chromosomes during synapsis in mice and humans (Yang et al. 2008). The *Tex11L* alternative splicing mutation exhibited a variable penetrance phenotype. All animals developed testicular phenotypes over time (decreased testis weight, abnormal seminiferous tubule morphology, and reduced sperm counts). One third of *Tex11L* mice had a prominent testicular and epididymal phenotype at 24, 36 and 52 weeks (3 of 9 mice) and infertility (2 of 6 mice).

There is an unusually high incidence of *TEX11* mutations discovered in men with idiopathic NOA. Yatsenko et. al. screened 289 men with NOA and 384 controls. *TEX11* mutations were found in 2.4% of the NOA patient cohort (Yatsenko et al. 2015). Similarly, Yang and colleagues screened 246 men with NOA and 175 controls and found *TEX11* mutations in 1% of the NOA cohort. *TEX11* is located on the X-chromosome and men, who have only one X chromosome, may be particularly susceptible to pathogenic variants in X-linked genes that are important for spermatogenesis. Furthermore, Wang and colleagues (Wang et al. 2001) reported that there is an abundance of X-linked genes expressed in spermatogonia. Using cDNA subtractive hybridization, the group identified 23 spermatogonia-expressed genes, 19 of which were novel and were named *testis-expressed* (*Tex*) genes. Of those 23 genes, 13 have human homologs, including TEX11; 10 are located on the X chromosome; and 3 are located on the Y-chromosome. Three *TEX* genes have been linked to infertility in the OMIM database (Online Mendelian Inheritance in Man): *TEX11, TEX14,* and *TEX15*. The human *TEX14* and *TEX15* genes are located on chromosomes 11 and 8, respectively. *TEX14* mutations have been identified in patients with NOA (Gershoni et al. 2017; An et al. 2021) and LoF mutations in mice cause azoospermia (Yang et al. 2008; Wang et al. 2021). Similarly, *TEX15* mutations have been identified in patients with unexplained NOA (Qureshi et al. 2023) and mouse knockouts were azoospermic (Yang et al. 2008; Qureshi et al. 2023).

Taken together, our study validated the association between the *TEX11D* (frameshift) and *TEX11L* (splicing) mutations and infertility. Mice with the human *TEX11A* (missense) mutation were fertile. As knowledge about validated NOA-associated genetic variants increases, this information could be used to develop diagnostic screens, and the value of those screens depends on the quality of evidence supporting the gene-disease relationships (Tüttelmann et al. 2018; Oud et al. 2019). Furthermore, diagnosis of single gene mutations that are strongly associated with NOA could identify targets for gene therapy. For example, Wang et. al. successfully corrected the *Tex11D* frameshift mutation (p.D435fs) in cultured mouse spermatogonial stem cells and restored spermatogenesis, leading to the successful production of offspring (Wang et al. 2021).

This same workflow could be used to correct monogenic causes of NOA in humans once robust methods to maintain and expand human SSCs in culture are established (David and Orwig 2020). Alternatively, gene editing could be performed in patient-derived induced pluripotent cells (iPSCs), which can be maintained, screened and expanded in long-term culture (Jang and Ye 2016). Gene-corrected iPSCs could then be differentiated into sperm, *in vivo* or *in vitro*, a feat that has been accomplished in mice (Hayashi et al. 2011; Ishikura et al. 2016) but not yet with human iPSCs.

Heritable human genome editing is controversial (Angrist et al. 2020) and currently illegal in the United States (H.R.2029). There is international consensus that human embryos that have been edited should not be used to create a pregnancy until safety for the unborn child and subsequent generations have been demonstrated. While the path to gene therapy for germline mutations like Tex11 is currently blocked, the British Royal Society, the US National Academy of Sciences and the US National Academy of Medicine published a consensus report outlining stringent preclinical and clinical requirements for establishing safety and efficacy that may provide a roadmap for clinical application in the future (Callaway 2016; National Academy of Medicine, National Academy of Sciences2020)

## Materials and methods

### Variant Selection

We screened NOA associated TEX11 variants reported by our group and others (Table 1) to identify potentially pathogenic frameshift, splice site, and missense mutations. I*n silico* analyses according to the American College of Medical Genetics standards were used to predict human variant pathogenicity. This missense variant (p.A683T, *Tex11A*), a variant of uncertain significance is in a currently uncharacterized region of the gene (Yatsenko et al. 2015). The splice site variant (p.L249, *Tex11L*) is a novel variant in the well conserved meiosis specific Spo22/ZIP4/TEX11 functional domain of the gene (Yatsenko et al. 2015). The frameshift variant (p.D435fs, *Tex11D*) features an insertion of TT in the TPR-like helical domain (Yang et al. 2015).

### Animals

Animals were maintained and housed in the laboratory animal facility at Magee-Womens Research Institute. All study procedures were approved by the Institutional Animal Care and Use Committee of the University of Pittsburgh and Magee-Womens Research Institute. Procedures were performed in accordance with the NIH Guide for the Care and Use of Laboratory Animals (Assurance # 3654-01).

### Generating Tex11 Mouse Models

For each of the *Tex11* variants, sgRNA sequences were designed to target near the variant location in the genome and were created with the program from Feng Zhang’s lab (https://zlab.bio/guide-design-resources)(Table 2). sgRNAs were produced in house using pSpCas9(BB)-2A-GFP (PX458) (Addgene 48138) as the plasmid DNA template (Table 2) and T7 in vitro transcription (MAXIscript™ T7 Transcription Kit, ThermoFisher), then purified using RNA Cleanup kit (Qiagen). The ssODNs were designed to knock in the human NOA-associated variant into the orthologous region of the mouse genome and were from Integrated DNA Technologies (Coralville, Iowa, Table 2).

B6D2F1 (C57BL/6J x DBA/2J F1) females (7-8 weeks old) were stimulated by intraperitoneal administration of 5 IU of PMSG (ProspecBio, Cat # HOR-272) at 3:30 pm on day 1 and 5 IU of hCG (Sigma-Aldrich, Cat # CG5-1VL) 48 hours later. Then the females were mated to B6D2 F1 males. The embryos were collected from oviducts the following morning. After the cumulus cells were removed by incubation in 1% hyaluronidase in M2 medium, the embryos were cultured in KSOM medium at 5% CO_2_, 37°C until electroporation.

The reagents for embryo electroporation were mixed as follows in Opti-MEM medium (Gibco, 31985070): Cas9 protein 100 ng/μL (Alt.R S.p. Cas9 Nuclease 3NLS IDT Cat # 1074181); Tex11 sgRNA 200 ng/μL, Tex11 ssODN 200 ng/μL. The electroporation was performed using the Super electroporator NEPA21 type IWE and CUY 501-1-1.5 electrode (NEPA GENE Co. Ltd, Chiba, Japan) with poring pulse (voltage 40 V, pulse length 2.5 ms, pulse interval 50 ms, number of pulses 4, decay rate 10%, polarity +); transfer pulse (voltage 7 V, pulse length 50 ms, pulse interval 50 ms, number of pulses 5, decay rate 40%, polarity +/-). After electroporation, the embryos were washed two times in KSOM medium, then cultured in KSOM medium overnight at 5% CO_2_, 37°C. On the following day, the two cell stage embryos were transferred to the oviducts of pseudopregnant CD1 females (0.5 dpc).

### Genotyping

Litters were genotyped at the time of weaning using DNA from ear clip biopsies. The biopsies were put in lysis buffer (5M NaCl, 0.5M EDTA, 1M Tris-HCl pH=8, 10% SDS, 1% β-Mercaptoethanol) and agitated at 55°C overnight. Samples were vortexed the next morning and spun at 11,000 x g for 2 min. The supernatant was transferred to a clean tube and 1 mL of 100% ethanol was added to the solution. Samples were vortexed and spun at 11,000 x g for 7 min. The supernatant was discarded and precipitated DNA pellets were dried in air for 1 hour. Pellets were resuspended in 50 mL of ultrapure, nuclease free H_2_O.

For *Tex11D*, PCR used LongAmp® Taq 2X Master Mix (M0287, New England Biolabs, MA, USA). *Tex11D* Fwd Primer will anneal to DNA of both *Tex11D* and wild-type mice. *Tex11D* Fwd wild-type primer will anneal with DNA from wild-type mice only. *Tex11D* Fwd Mutant Primer will anneal to DNA from *Tex11D* mutant mice only. *Tex11D* Rev Primer was used for all PCR reactions (Suppl. Fig. 1). The PCR cycle was 95°C for 10 minutes then 95°C melting temperature (30s), annealing temperature (55°C Fwd and Fwd wild type; 57°C Fwd mutant) (20s) and 72°C extension temperature (50s) for 30 cycles, then 72°C for 10 minutes. PCR products were run on a 1.5% agarose gel at 110 volts for 15 minutes and imaged for bands.

For the *Tex11A* and *Tex11L*, PCR was performed using master mix prepared with components from Kapa Biosystems, Inc. (Wilmington, MA, USA): 5 uL HiFi Fidelity Buffer 5X, 1 uL 10 mM KAPA dNTP, 15.5 uL nuclease-free water, and 0.5 uL 1 U/uL KAPA HotStart enzyme combined with 0.5 uL of 10 uM of both forward and reverse primers. Forward and reverse primers for both the *Tex11A* and *Tex11L* lines (Suppl. Fig. 1). The PCR cycle for the A line was 95°C for 5 minutes, then 98°C melting temperature (20s), 64°C annealing temperature (25s), and 72°C extension temperature (25s) for 39 cycles, then 72°C for 5 minutes. For the *Tex11L* line, the PCR cycles the annealing temperature was 60°C. The PCR samples were then run on a 1.3% agarose gel at 90 V and 400 mA for 50 minutes and imaged for bands. The PCR products underwent Sanger sequencing performed by Eton Bioscience (San Diego, CA, USA) to determine genotype.

Breeding colonies for each line were established by breeding mutant founders to DBA/2J mice to produce experimental knock-in mice and littermate controls.

### Standard Breeding Fertility Analysis

At 2 months old, hemizygous males (Tex11^-/Y^) from *Tex11D, Tex11L, Tex11A,* and wild type littermate controls were bred with two wild-type female for two weeks. After a one-week break, males were bred with two new wild-type females. After breeding trials, mice were sacrificed, and testes and epididymides were collected for further analysis.

### Long-Term Breeding studies

To assess long-term effects of the *Tex11L* splice variant, 3 hemizygous mutant and 3 wild-type controls were bred with wild-type and heterozygous females from the respective lines, swapping every two weeks for 52 weeks. At the end, mice were sacrificed and samples were collected as described above.

### Immunohistochemistry

Mouse testis and epididymal tissues were removed from three *Tex11D*, *Tex11A*, and *Tex11L* mutant mice and three wild-type control mice between 8-12 weeks of age. The wild-type control mice and *Tex11D* tissues were fixed in 4% paraformaldehyde in Dulbecco’s phosphate-buffered saline (DPBS) overnight at 4°C. The *Tex11A* and *Tex11L* mutant mice were fixed in 10% formalin overnight at 4°C. Tissues were washed with room temperature DPBS three to four times with at least 60-minute intervals between washes before processing for paraffin embedding. Tissue sections (0.5 µm) were mounted on glass slides for immunohistochemistry.

Sections were warmed on a slide warmer for 10 minutes and then deparaffinized with xylene (3 x 10 minutes). They were then hydrated in graded ethanol with two changes of 100% ethanol, followed by 95%, 80%, 70%, 50%, and 25% ethanol each for 5 minutes. The slides were rinsed once in 1X dPBS and once in dH_2_O and immersed in 97.5°C Tris antigen retrieval buffer solution for 40 minutes. After a cooling period of 20 minutes, the slides were rinsed in Phosphate Buffered Saline with Tween-20 (PBS-T) twice for two minutes each. Tissue sections were then permeabilized on a shaker for 30 minutes in a solution containing 100 mL 0.02% Triton X in 1x PBS. After two subsequent washes with PBS-T, sections were incubated with a blocking buffer (10% normal donkey serum, 3% BSA Fraction IV, 0.2% Triton X-100, 0.3M Glycine, 1X dPBS-T) for two hours at room temperature in a humidified chamber.

The following primary antibodies, diluted in blocking buffer, were used for immunofluorescence staining of the mouse testis tissue sections: rabbit anti-SALL4 (Abcam, diluted 1:800), rabbit anti-STRA8 (Abcam, diluted 1:200), goat anti-PIWIL1 (R&D Systems, diluted 1:200), goat anti-TP1 (Abcam, diluted 1:1,500), goat anti-ZIP4H/Tex11 (R&D Systems, diluted 1:1,000), goat anti-VASA (R&D Systems, diluted 1:200). These primary antibodies were incubated with tissue sections overnight at 4°C in a humidified chamber. Isotype matched normal IgG controls were used on one tissue section on each slide instead of primary antibody as a negative control. After three two-minute washes with PBS-T, secondary donkey anti-rabbit (Invitrogen, diluted 1:200) or donkey anti-goat AlexaFluor-488 (Thermo Fisher, diluted 1:200) with donkey anti-goat AlexaFluor-568 conjugated IgG (Invitrogen, diluted 1:200 in blocking buffer) were applied to the appropriate tissue sections and incubated at room temperature for two hours in a humidified chamber. Sections were washed with PBS-T three times for two minutes each and incubated in a humidified chamber for 15 minutes at room temperature with Vectashield medium with 4’,6-diamidino-2-phenylindole (DAPI) (Vector Labs) for fluorescence imaging. After two final washes for two minutes each in deionized water, the slides were mounted with Vectashield medium. Imaging was performed on the Nikon Eclipse 90i fluorescence microscope, using an X-Cite 120 fluorescence light source, Andor Zyla fluorescent camera, and Nikon Eclipse 90i software.

### Meiotic Spreads

Prophase I spermatocyte chromosome spreads were performed as previously described (Brieño-Enríquez et al. 2016) in 4 males per genotype. Primary antibodies included: anti-mouse SYCP3 (ab97672), SYCP1 (ab15090), yH2AX (ab26350), RPA (Cell Signaling #2208S), RAD51 (Millipore #PC130-100U) and MLH1 (BD Biosciences Pharmingen, #550838). Alexa Fluor secondary antibodies (Jackson Immunoresearch) were used. Image acquisition was performed using a Zeiss AxioImager M2. Microscope Images were processed using Zeiss efficient navigation (ZEN) software.

### Sperm analysis

To assess sperm production, the two caput epididymides and one cauda epididymis were fixed in 4% PFA or 10% formalin for histological analysis and one cauda epididymis from each mouse was dissected to collect sperm. To collect sperm, a 200 µL drop of sterile, warm PBS was placed in the center of a two-centimeter dish under mineral oil. The cauda epididymis was placed in the PBS droplet after it was cleaned of fat and extraneous tissue, minced with scissors to release the sperm, and then incubated at 37°C for 15 minutes.

The sperm droplet was then aspirated, added to an Eppendorf tube and gently pipetted. Ten microliters of the sperm solution were diluted (50-fold) into 490 µL of warm PBS in a new Eppendorf tube to determine count and viability using a hemacytometer.

### Statistical Analysis

A two-sample proportion test with continuity correction was performed to compare the proportion of matings produced between wild type and mutant mouse lines. The p-value was adjusted for multiple comparisons using Holm’s method. The rest of the data was log transformed and a small offset value of 0.01 was used to data that contained a 0. A two-sample t-test with unequal variances adjusted for multiple comparisons using Holm’s method were performed on the remaining data. Tubule area was analyzed by transforming the data to a log scale and then using a t-test the area between wild type and mutant was performed. For meiotic spread data, RAD51 and RPA foci were compared using an ordinary one-way ANOVA adjusted for multiple comparisons and MLH1 were compared with an unpaired t-test.

## Acknowledgements

This study was supported by the *Eunice Kennedy Shriver* National Institute for Child Health and Human Development (NICHD) grant HD096723 to ANY and KEO. GRA was supported by NICHD grant HD114406. We acknowledge the contributions of the Genome Editing, Transgenic and Virus (GETV) Core, the Histology Core of Magee-Womens Research Institute, and Jannah Kuong for their help with data generation and analysis.

GRA, RHG, CA, ANY, and KEO designed experiments. GRA, RHG, CA, CP, EB, JJH, MS, NP, and MM conducted experiments and collected data. YS performed CRISPR/Cas9 edits, ICSI, and embryo transfers. CC analyzed histological data. TC conducted statistical analysis. GRA, RHG, and CA wrote the manuscript. GRA and KEO revised the manuscript. All authors contributed to the article and approved the submitted version.

---

**Supplemental Figure 1.**
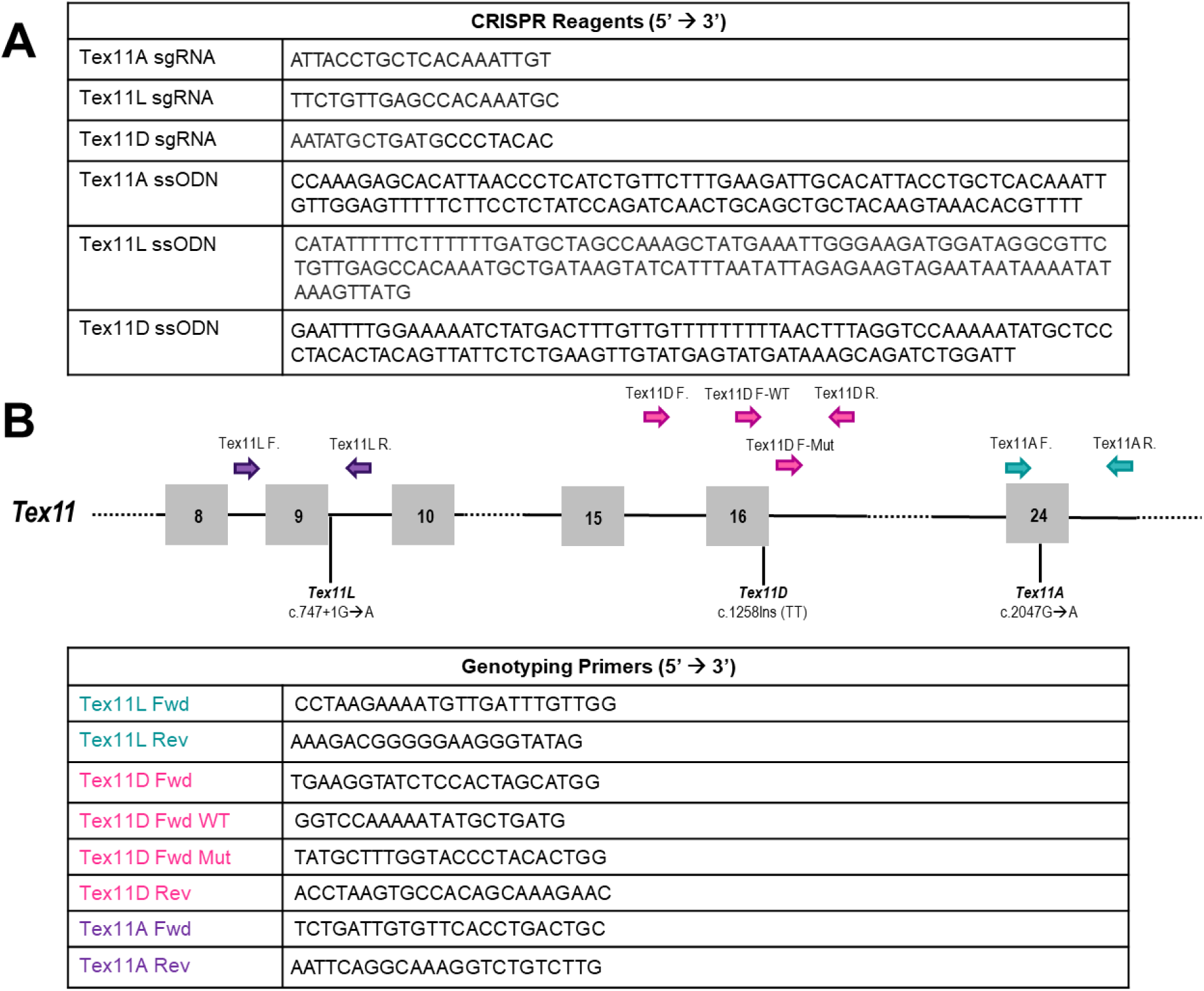
CRISPR/Cas9 Knock-in Mouse Model Reagents (*A*) Single guide RNA and ssODN sequences used to generate knock-in of human TEX11 variants into the orthologous region of the mouse genome. (*B*) Schematic of Tex11 gene locus demonstrating the location of the genotyping primers used for each line in relation to variant. Grey numbered squares represent exons and black lines represent introns.

**Supplemental Figure 2.**
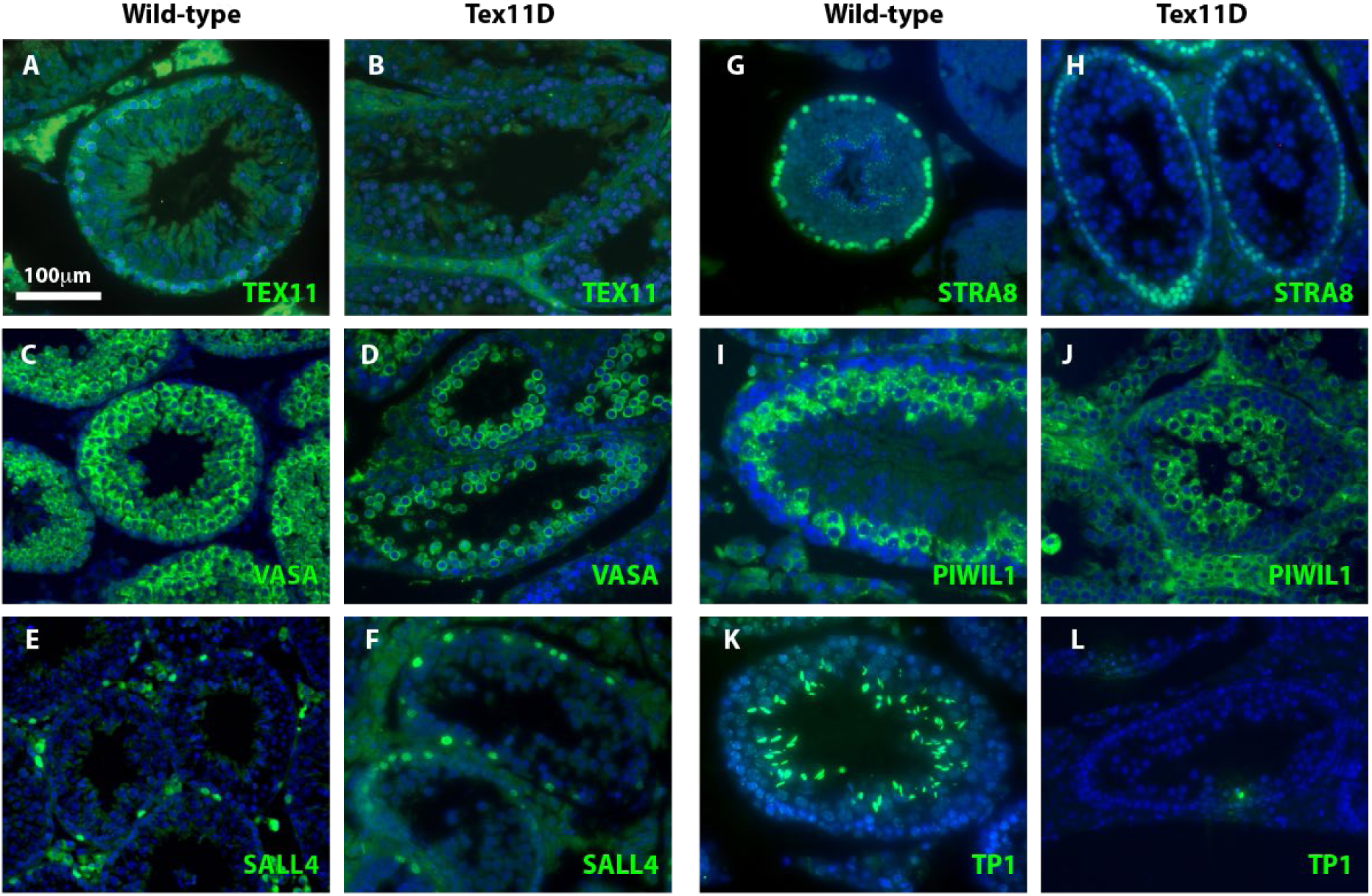
T*e*x11D mice have a meiotic arrest phenotype. (*A, B*) Wild type (WT) show expression of TEX11 in spermatocyte while *Tex11D* mice have no TEX11 protein expression. (*C, D*) *Tex11D* mice have a reduced number of VASA+ germ cells compared to WT mice. (*E-J*) Both WT and *Tex11D* mice had SALL4+ undifferentiated spermatogonia, STRA8+ differentiating spermatogonia and PIWIL1+ spermatocytes. (*K, L*) Wild type mice had TP1+ spermatids while *Tex11D* mice had no TP1+ spermatids. These demonstrate that *Tex11D* mice had a meiotic arrest at the spermatocyte stage of spermatogenesis. Scale bar in all images is 100 µm.

**Supplemental Figure 3.**
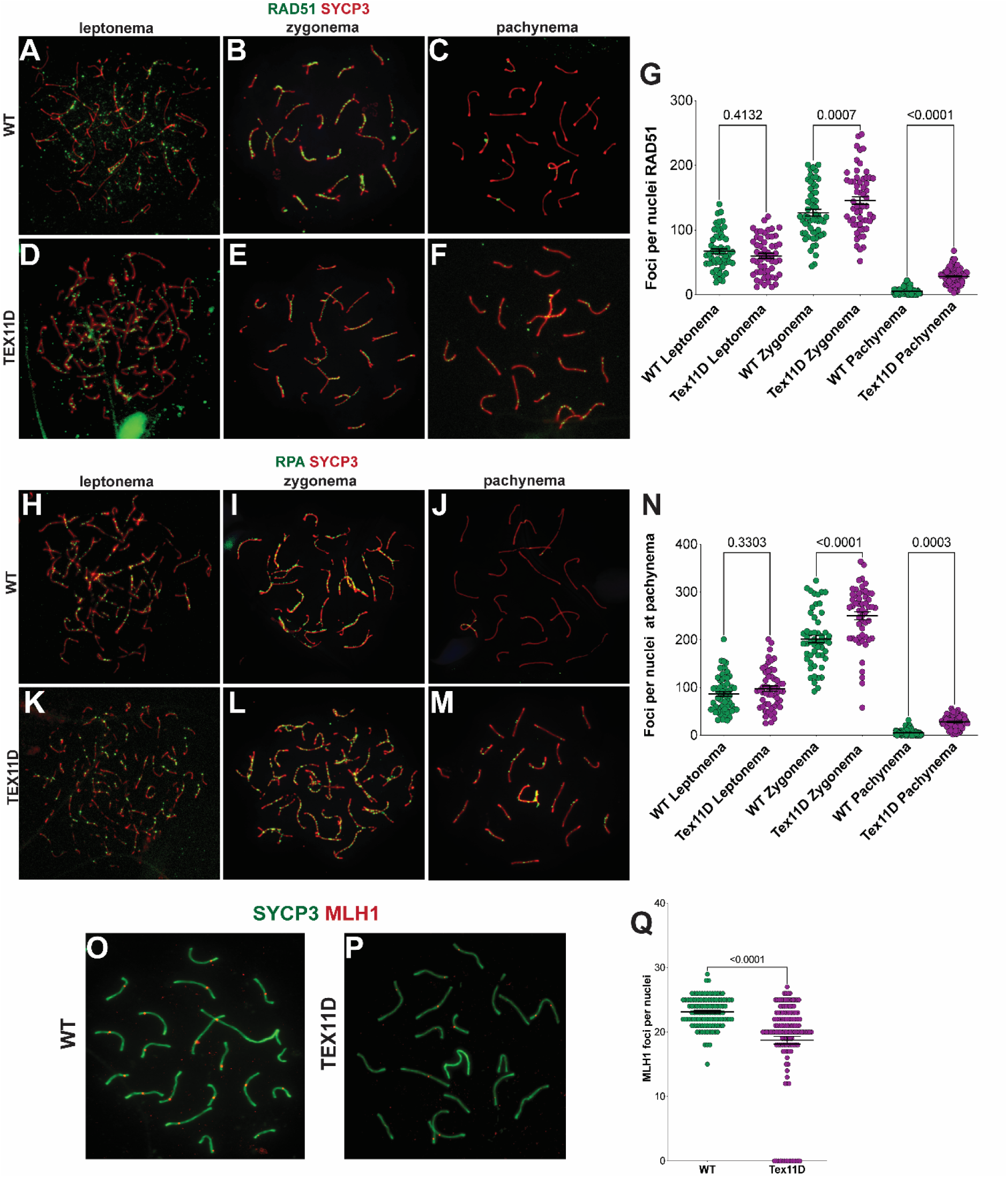
T*e*x11D mice exhibit unresolved DNA crossover events. (*A-F*) Meiotic spreads staining for SYCP3 (chromosomes, red) and RAD51 (double strand break foci green) through leptonema, zygonema and pachynema stages in prophase I in wild type (WT, *A-C*) and *Tex11D* (D-F) are quantified in (*G*). (*H-M*) SYCP3 (red) and RPA (single strand DNA, green) in WT (*H-J*) and *Tex11D* (*K-M*) are quantified in (*N*). (*O, P*) Meiotic spreads staining for SYCP3 (chromosomes, green) and MLH1 foci (DNA crossovers, red) in WT (*O*) and Tex11D (*P*) are quantified in (*Q*).

**Supplemental Figure 4.**
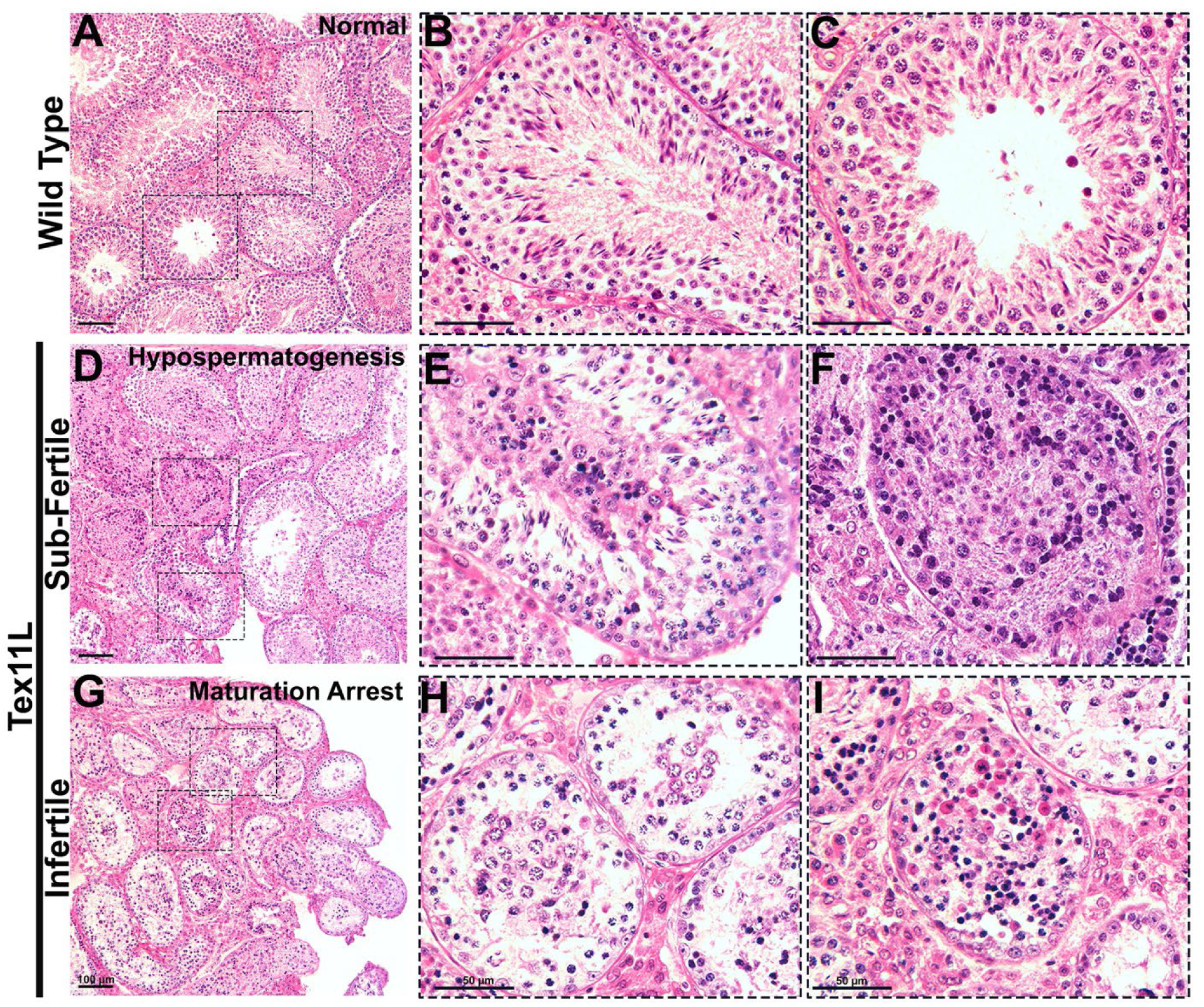
T*e*x11L mice have abnormal seminiferous tubule morphology at 52 weeks. (*A*) Hematoxylin and eosin (H&E) stain of testicular seminiferous tubules at 52 weeks of age for wild type (WT) mice are magnified in (*B*, *C*). H&E stain of *Tex11L* sub-fertile mouse seminiferous tubules are magnified in (*E, F*). H&E stain of *Tex11L* infertile mouse seminiferous tubules are magnified in (*H, I*), indicating disorganization of germ cells, maturation arrest, and tubule atrophy. Scale bar is 100 µm for (*A, D and G*) and 50 µm for magnified images (*B, C, E, F, H, I*).

